# Footprints of antigen processing boost MHC class II natural ligand predictions

**DOI:** 10.1101/285767

**Authors:** Carolina Barra, Bruno Alvarez, Sinu Paul, Alessandro Sette, Bjoern Peters, Søren Buus, Massimo Andreatta, Morten Nielsen

## Abstract

**Background:** Major Histocompatibility complex class II (MHC-II) molecules present peptide fragments to T cells for immune recognition. Current predictors for peptide:MHC-II binding are trained on binding affinity data, generated *in-vitro* and therefore lacking information about antigen processing.

**Methods:** We generate prediction models of peptide:MHC-II binding trained with naturally eluted ligands derived from mass spectrometry in addition to peptide binding affinity datasets.

**Results:** We show that integrated prediction models incorporate identifiable rules of antigen processing. In fact, we observed detectable signals of protease cleavage at defined positions of the ligands. We also hypothesize a role of the length of the terminal ligand protrusions for trimming the peptide to the MHC presented ligand.

**Conclusions:** The results of integrating binding affinity and eluted ligand data in a combined model demonstrate improved performance for the prediction of MHC-II ligands and T cell epitopes, and foreshadow a new generation of improved peptide:MHC-II prediction tools accounting for the plurality of factors that determine natural presentation of antigens.

## Background

Major Histocompatibility Complex Class II (MHC-II) molecules play a central role in the immune system of vertebrates. MHC-II present exogenous, digested peptide fragments on the surface of antigen presenting cells, forming peptide-MHC-II complexes (pMHCII). On the cell surface, these pMHCII complexes are scrutinized, and if certain stimulatory conditions are met, a T helper lymphocyte may recognize the pMHCII and initiate an immune response (Rudolph, Stanfield, & Wilson, 2006).

The precise rules of MHC class II antigen presentation are influenced by many factors including internalization and digestion of extracellular proteins, the peptide binding motif specific for each MHC class II molecule, and the transport and surface half-life of the pMHCIIs. The MHC-II binding groove, unlike MHC class I, is open at both ends. This attribute facilitates peptide protrusion out of the groove, thereby allowing longer peptides (and potentially whole proteins) to be loaded onto MHC-II molecules (Kim et al., 2014; Sette, Adorini, Colon, Buus, & Grey, 1989). Peptide binding to MHC-II is mainly determined by interactions within the peptide binding groove, which most commonly encompass a peptide with a consecutive stretch of 9 amino acids (Andreatta, Jurtz, et al., 2017). Ligand residues protruding from either side of the MHC binding groove are commonly known as peptide flanking regions (PFRs). The PFRs are variable in length and composition, and affect both the peptide MHC-II binding (Lovitch, Pu, & Unanue, 2006), and the subsequent interaction with T cells (Arnold et al., 2002; Carson, Vignali, Woodland, & Vignali, 1997; Godkin et al., 2001). The open characteristic of the MHC-II binding groove does not constrain the peptides to a certain length, thereby increasing the diversity of sequences that a given MHC-II molecule can present. Also, MHC-II molecules are highly polymorphic, and their binding motifs have appeared to be more degenerate than MHC-I motifs (Andreatta, Schafer-Nielsen, Lund, Buus, & Nielsen, 2011; Hammer et al., 1993; Sturniolo et al., 1999).

Considering all the aspects mentioned above, MHC-II motif characterization and rational identification of MHC-II ligands and epitopes is a highly challenging and costly endeavor. Because MHC-II is a crucial player in the exogenous antigen presentation pathway, considerable efforts have been dedicated in the past to develop efficient experimental techniques for MHC-II peptide binding quantification. The traditional approach to quantify peptide MHC-II binding relies on measuring binding affinity, either as the dissociation constant Kd of the complex (Roche & Cresswell, 1990; Hall et al., 2002), or in terms of IC50 (concentration of the query peptide which displaces 50% of a bound reference peptide) (Buus, Sette, Colon, Miles, & Grey, 1987). To date, data repositories such as the Immune Epitope Database (IEDB) (Vita et al., 2015) have collected more than 150,000 measurements of peptide-MHC-II binding interactions. Such data have been used during the last decades to develop several prediction methods with the ability to predict binding affinities to the different alleles of MHC class II. While the accuracy of these predictors has increased substantially over the last decades due the development of novel machine learning frameworks and a growing amount of peptide binding data being available for training (Andreatta, Trolle, et al., 2017), state-of-the-art methods still fail to accurately predict accurately MHC class II ligands and T cell epitopes (Gowthaman & Agrewala, 2008; Wang et al., 2008).

Recent technological advances in the field of mass spectrometry (MS) have enabled the development of high-throughput assays, which in a single experiment can identify several thousands of peptides eluted of MHC molecules (reviewed in (Caron et al., 2015)). Large data sets of such naturally presented peptides have been beneficial to define more accurately the rules of peptide-MHC binding (Abelin et al., 2017; Bassani-Sternberg & Gfeller, 2016; Bergseng et al., 2015; Chong et al., 2017; Clement et al., 2016; Jurtz et al., 2017; Ooi et al., 2017). For several reasons, analysis and interpretation of MS eluted ligand data is not a trivial task. Firstly, because any given individual constitutively expresses multiple allelic variants of MHC molecules; thus, the ligands detected by MS are normally a mixture of specificities, each corresponding to a different MHC molecule. Secondly, MHC-II ligands can vary widely in length, and identification of the binding motifs requires a sequence alignment over a minimal binding core. Finally, data sets of MS ligands often contain contaminants and false spectrum-peptide identifications, which add a component of noise to the data. We have earlier proposed a method capable of dealing with all these issues, allowing the characterization of binding motifs and the assignment of probable MHC restrictions to individual peptides in such MS ligand data sets (Andreatta, Alvarez, & Nielsen, 2017; Andreatta, Lund, & Nielsen, 2013).

Because naturally eluted ligands incorporate information about properties of antigen presentation beyond what is obtained from *in-vitro* binding affinity measurements, large MS-derived sets of peptides can be used to generate more accurate prediction models of MHC antigen presentation (Abelin et al., 2017; Bassani-Sternberg & Gfeller, 2016; Jurtz et al., 2017). As shown recently, generic machine learning tools, such as NNAlign (Andreatta et al., 2011; Nielsen & Andreatta, 2017), can be readily applied to individual MS data sets, which in turn can be employed for further downstream analyses of the immunopeptidome (Alvarez, Barra, Nielsen, & Andreatta, 2018) The amount of MHC molecules characterized by MS eluted ligand data is, however, still limited. This has led us to suggest a machine learning framework where peptide binding data of both MS and *in vitro* binding assays are merged in the training of the prediction method (Jurtz et al., 2017). This approach has proven highly powerful for MHC class I, but has not, to the best of our knowledge, been applied to MHC class II.

Undoubtedly, antigen processing plays a critical role in generating CD4+ T cell epitopes presented by MHC class II molecules. It is assumed that endo- and exo-peptidase activities, both before and after binding to the MHC-II molecule, play a key role in the generation and trimming of MHC class II ligands (Blum, Wearsch, & Cresswell, 2013; Lippolis et al., 2002). However, the precise rules of MHC class II antigen processing are poorly understood. Earlier works identified patterns of protein cleavage in HLA-DR ligands; Kropshofer et al. found proline at the penultimate N and C terminal position (Kropshofer et al., 1993) Ciudad et al. observed aspartic acid before the cleavage site and proline next to the cut sites in HLA-DR ligands (Ciudad et al., 2017). In contrast, Bird et al. suggested that endolysosomal proteases have a minor and redundant role in peptide selection leading to the conclusion that the effect of processing on the generation of antigenic peptides is “relatively non-specific” (Bird, Trapani, & Villadangos, 2009). Given this context, it is perhaps not surprising that limited work has been aimed at integrating processing signals into a prediction framework for MHC-II ligands.

In this work, we have analyzed large data sets of MS MHC-II eluted ligands obtained from different research laboratories covering three HLA-DR molecules with the purpose of investigating the consistency in the data; quantifying the differences in binding motifs contained with such MS eluted data compared to traditional *in vitro* binding data; defining a new machine-learning framework capable of integrating information from MS eluted ligand and *in vitro* binding data into a prediction model for MHC-II peptide interaction prediction; and finally evaluating if inclusion of potential signals from antigen processing are consistent between different data sets and can be used to boost the performance of peptide-MHCII prediction models.

## Methods

### Datasets

HLA class-II peptidome data were obtained from two recent MS studies. Three datasets corresponding to the HLA-DRB1:0101: DR1-Ph, DR1-Pm (Ooi et al., 2017), and DR1-Sm (Clement et al., 2016), two to DRB1*15:01: DR15-Ph and DR15-Pm, and one to the allele DRB5*01:01: DR51 Ph (for details see table 1). Here, the data sets with subscript h correspond to data obtained from human cell lines and data sets with the subscript m to data obtained from human MHC-II molecules transfected into MHC-II deficient mice cell lines. Details on how the data were generated is provided in the original publications. Note that DR15 Ph and DR51 Ph data sets were obtained from a heterozygous EBV transformed B lymphoblastoid cell line (BLCL), IHW09013 (also known as SCHU), which expresses two HLA-DR molecules, HLA-DRB1*15:01 and HLA-DRB5*01:01 (shortened here with the name DR15/51). The DR1 Ph dataset was extracted from a BLCL culture as well (IHW09004). On the other hand, DR1 Pm, DR1 Sm and DR15 Pm datasets were extracted from HLA transgenic mice, and therefore only cover the human alleles of interest. These cells are treated here as monoallelic.

**Table 1.**
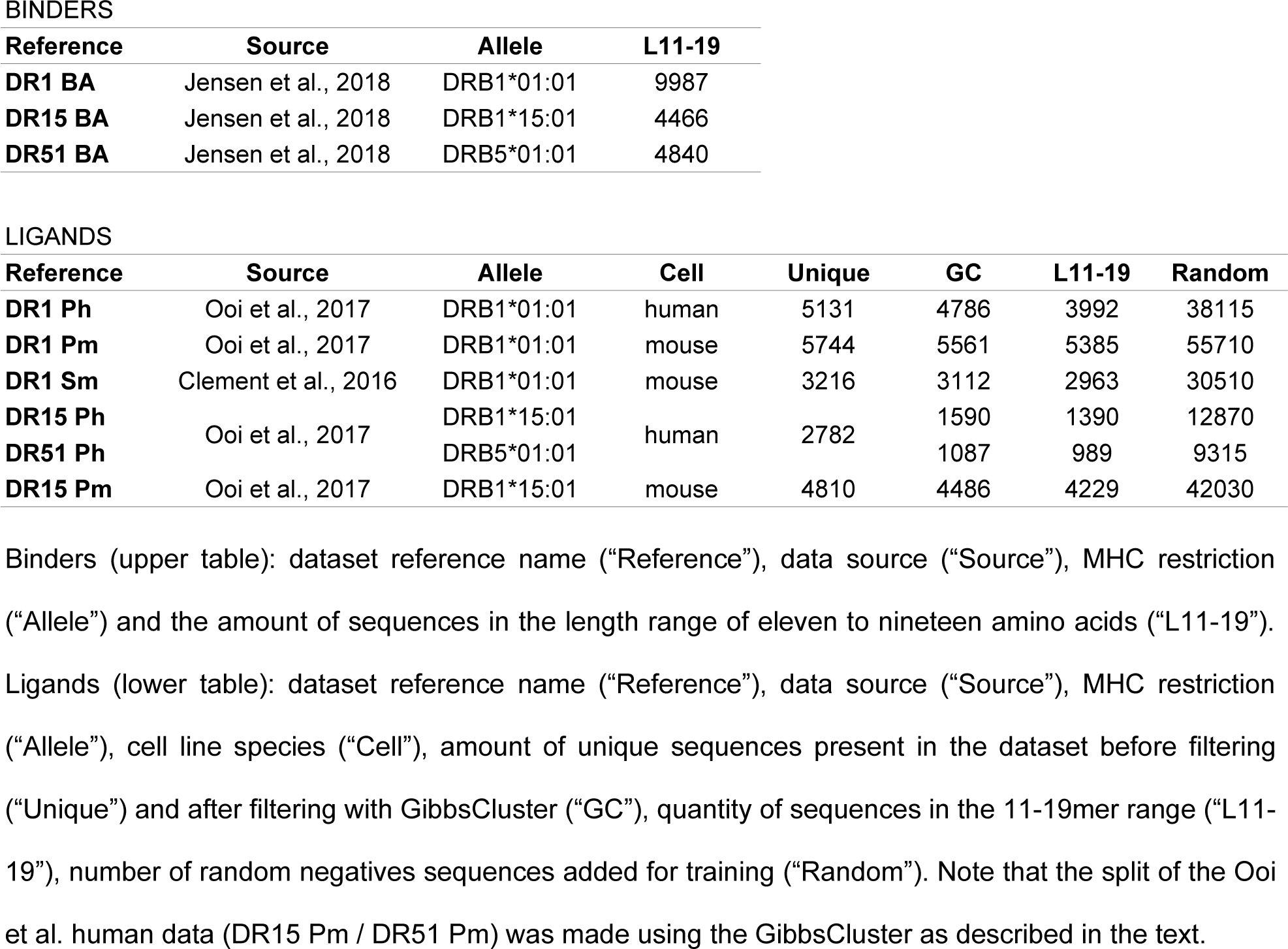
Summary of binding affinity (“Binders”) and eluted ligand (“Ligands”) datasets used in this work.

MHC class II peptide binding affinity data was obtained from previous publications (Jensen et al., 2018) for the alleles DR1 (DRB1*01:01, 9987 peptides) and DR15 (DRB1*15:01, 4,466 peptides), and DR51 (DRB5*01:01, 4840 peptides).

The MS-derived ligand datasets were filtered using the GibbsCluster-2.0 method with default settings as described earlier (Alvarez et al., 2018), to remove potential noise and biases imposed by some data containing multiple binding specificities. The details of the binding affinity (BA) and eluted ligand (EL) data sets are described in table 1.

### NNAlign modeling and architecture

Models predicting peptide-MHC interactions were trained as described earlier using NNAlign (Alvarez et al., 2018; Nielsen & Andreatta, 2017). Only ligands of length 11-19 amino acids were included in the training data. Random peptides of variable lengths derived from the non-redundant UniProt database were used as negatives. The same amount of random negatives was used for each length (11 to 19), and consisted of five times the amount of peptides for the most represented length in the positive ligand dataset. Positive instances were labeled with a target value of 1, and negatives with a target value of 0. Prior to training, the datasets were clustered using the common motif approach described earlier (Hobohm, Scharf, Schneider, & Sander, 1992) with a motif length of 9 amino acids to generate five partitions for cross-validation.

Two types of model were trained; one with single data type (eluted ligand or binding affinity) input, and one with a mixed input of the two data types. Single models per each dataset and allele were trained as previously described with either binding affinity or eluted ligand data as input (Alvarez et al., 2018). All models were built as an ensemble of 250 individual networks generated with 10 different seeds; 2, 10, 20, 40, and 60 hidden neurons; and 5 partitions for cross-validation. Models were trained for 400 iterations, without use of early stopping. Additional settings in the architecture of the network were used as previously described for MHC class II (Alvarez et al., 2018). Combined models were trained as described earlier (Jurtz et al., 2017) with both binding affinity and eluted ligand data as input. Training was performed in a balanced way so that on average the same number of data points of each data type (binding affinity or eluted ligand) are used for training in each training iteration.

Novel modifications were introduced to the architecture of NNAlign to better account for specific challenges associated with MHC class II ligand data. For the network to be able to learn peptide length preferences, a “binned” encoding of the peptide length was introduced, consisting of a one-hot input vector of size nine (one neuron for each of the lengths 11 to 19). In order to guide binding core identification, a burn-in period was introduced with a limited search space for the P1 binding core position. During the burn-in period, consisting of a single learning iteration, only hydrophobic residues were allowed at the P1 binding core anchor position. Starting from the second iteration, all amino acids were allowed at the P1 position (Supplementary figure 1a and 1b).

### NetMHCII and NetMHCIIpan

NetMHCII version 2.3 (Jensen et al., 2018) and NetMHCIIpan version 3.2 (Jensen et al., 2018), peptide:MHC-II binding affinity prediction algorithms were employed in this work as a benchmark comparison for the new proposed model.

### Sequence logos

Sequence logos for binding motifs were constructed using Seg2Logo tool with default parameters (Thomsen & Nielsen, 2012). Logos for context information were generated as weighted Kulback-Leibler logos using Seq2Logo excluding sequence weighting. Amino acids were grouped by negatively charged (red), positively charged (blue), polar (green) or hydrophobic (black).

### Performance Metrics

In order to assess the performance of our new model, we employed three different and well-known metrics: AUC (Area Under the ROC Curve); AUC 0.1 (Area Under the ROC Curve integrated up to a False Positive Rate of 10%); and PPV (Positive Predictive Value). AUC is a common performance measurement for predictive models, which takes into account the relationship between True Positive Rates (TPR) and False Positive Rates (FPR) for different prediction thresholds. AUC0.1 is similar to AUC but focuses on the high specificity range of the ROC curve. PPV is here calculated by sorting all predictions and estimating the fraction of true positives with the top N predictions, where N is the number of positives in the benchmark data set. PPV represents a good metric to benchmark on highly unbalanced datasets like MS-derived elution data, where we have approximately ten times more negatives than positives.

## Results

### Data filtering and motif deconvolution

We first set out to analyze the different MS data sets of eluted ligands. Data were obtained from two recent publications; Ooi et al. (Ooi et al., 2017) (termed P) and Clement et al. (Clement et al., 2016) (termed S) covering the HLA-DRB1*01:01, HLA-DRB1*15:01 and HLA-DRB5*01:01 MHC class II molecules. Data were obtained from either human (termed h) or HLA-DR transfected mice (termed m) cell lines. Using this syntax, DR1 Ph corresponds to the HLA-DRB1*01:01 data from the human cell in the study by Ooi et al. (for more details see Materials and Methods). Here, we applied the GibbsCluster method with default parameters for MHC class II to both filter out potential noise and to identify the binding motif(s) contained in each data set. The result of this analysis is shown in Figure 1, and confirms the high quality of the different ligand data sets. In all data sets less than 7% of the peptides were identified as noise (assigned to the trash cluster), and in all cases GibbsCluster did find a solution with a number of clusters matching the number of distinct MHC specificities present in a given data set. In this context, the DR15 Ph is of particular interest, since this data set was obtained from a heterozygous cell line expressing two HLA-DR molecules, HLA-DRB1*15:01 and HLA-DRB5*01:01 (shortened here as DR15/51 Ph). Consequently, this data set contains a mixture of peptides eluted from both of these HLA-DR molecules. The GibbsCluster method was able to handle this mixed data set, and correctly identified two clusters with distinct amino acid preferences at the anchor positions P1, P4, P6 and P9. Moreover, a comparison of the motifs identified from the different data sets sharing the exact same HLA-DR molecules revealed a very high degree of overlap, again supporting the high accuracy of both the MS eluted ligand data and of the GibbsCluster analysis tool.

**Figure 1.**
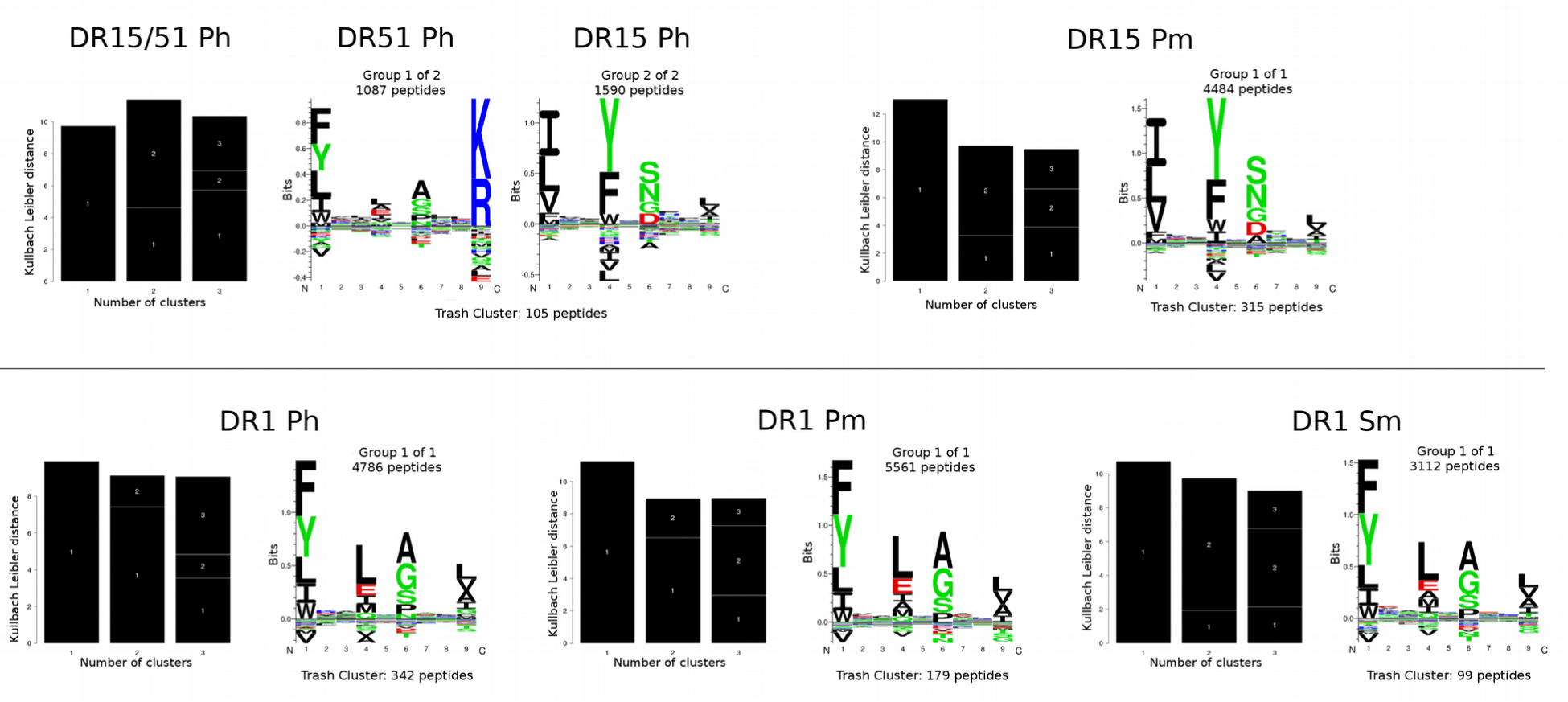
GibbsCluster output for the five eluted ligand datasets employed in this work. For each set, the Kullback-Leibler distance (KLD) histogram (black bars) is displayed, which indicates the information content present in all clustering solutions (in this case, groups of one to three clusters) together with the motif logo(s) corresponding to the maximum KLD solution. The upper row gives the results for the DR15/51 datasets; the lower row for the DR1 datasets. Note that DR15 Ph was obtained from a cell line which expresses two HLA-DR molecules, HLA-DRB1*15:01 and HLA-DRB5*01:01 (DR15/51).

### Training prediction models on MHC class II ligand data

After filtering and deconvolution with GibbsCluster, MHC peptide binding prediction models were constructed for each of the 6 data sets corresponding to the majority clusters in Figure 1. Models were trained using the NNAlign framework as described in materials and methods. The eluted ligand data sets (EL) were enriched with random natural peptides labeled as negatives, as described in Material and Methods. Likewise, models were trained and evaluated on relevant and existing data sets of peptide binding affinities (BA) obtained from the IEDB (Jensen et al., 2018; Vita et al., 2015), as described in Material and Methods. These analyses revealed a consistent and high performance for the models trained on the different eluted ligand data sets (Table 2). In accordance with what has been observed earlier for MHC class I (Jurtz et al., 2017), the overall cross-validated performance of models trained on binding affinity data is lower than that of models trained on eluted ligand data. Note that this observation is expected due the very different nature of the binding affinity and eluted ligand data sets: eluted ligand data are highly unbalanced, categorized, and prefiltered to remove ligands not matching the consensus binding motif.

**Table 2.** Cross-validation performance of models trained on Binding Affinity (BA) or Eluted Ligands (EL) data.

| BA Single Models |  |  | EL Single Models |  |  |  |
| --- | --- | --- | --- | --- | --- | --- |
| Training set | AUC | AUC 0.1 | Training set | AUC | AUC 0.1 | PPV |
| DR1 BA | 0.84 | 0.374 | DR1 Ph | 0.96 | 0.819 | 0.773 |
|  |  |  | DR1 Pm | 0.986 | 0.888 | 0.815 |
|  |  |  | DR1 Sm | 0.977 | 0.851 | 0.794 |
| DR15 BA | 0.843 | 0.287 | DR15 Ph | 0.987 | 0.901 | 0.85 |
|  |  |  | DR15 Pm | 0.989 | 0.917 | 0.859 |
| DR51 BA | 0.846 | 0.38 | DR51 Ph | 0.961 | 0.759 | 0.717 |

The binding motifs captured by the different models are shown in Figure 2. As evidenced by identical anchor positions (P1, P4, P6 and P9) and virtually identical anchor residues, highly consistent motifs were obtained from the same HLA-DR molecules irrespective of the source of the peptide (i.e. whether they were obtained from human or mouse cells, or from different laboratories). This observation to a high degree extended to the motifs obtained from binding affinity data, although we did observe subtle, but consistent, differences between the binding motifs derived from eluted ligand and peptide binding affinity data, exemplified for instance by the preference for E at P4 and for D at P6 in the eluted ligand motifs for DR1 and DR15, respectively. Such preferences are absent from the motifs derived from the peptide binding affinity data.

**Figure 2.**
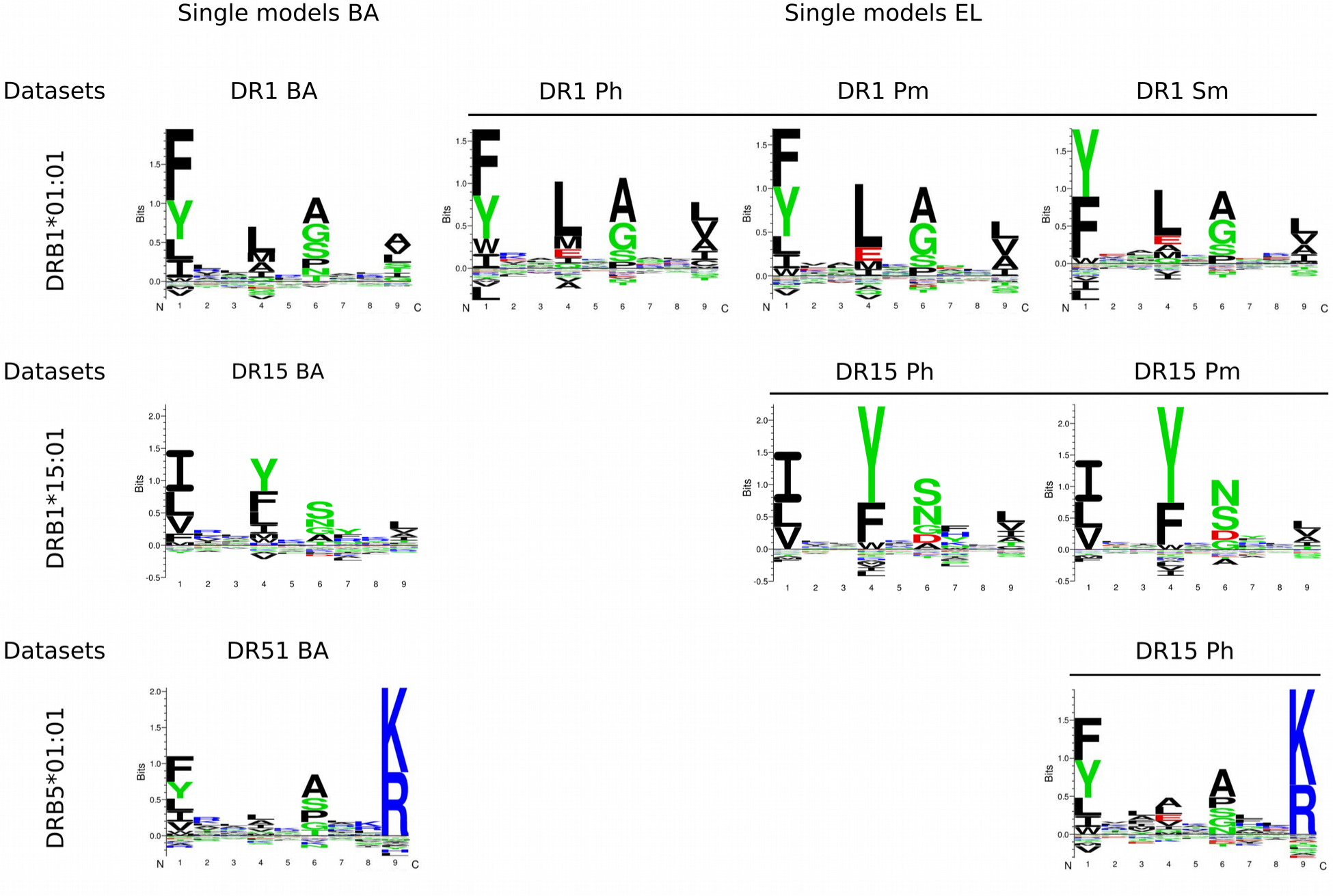
Binding preferences learned by the single NNAlign (Nielsen & Andreatta, 2017) models trained on Binding Affinity (BA) or Eluted Ligands (EL) data. In the top row, motifs for the DRB1*01:01 allele are shown, with overlined logo-plots (right) corresponding to models trained on EL data, and the non-overlined logo (left) corresponding to the BA trained model. Similarly, binding motifs for DRB1*15:01 are displayed in the bottom row, with overlined logos (right) also indicating the EL-trained models preferences, and the non-overlined logo-plot (left) indicating the BA preference. Logos were constructed from the predicted binding cores in the top 1% scoring predictions of 900.000 random natural peptides for BA and from the top 0.1% scoring predictions for EL.

### Training a combined prediction model on MHC-II binding affinity and ligand elution data

Earlier work on MHC class I has demonstrated that the information contained within eluted ligand and peptide binding affinity data is, to some degree, complementary, and that a prediction model can benefit from being trained integrating both data types (Jurtz et al., 2017). Here, we investigate if a similar observation could be made for MHC class II. As proposed by Jurtz et al., we extended the NNAlign neural network model to handle peptides from both binding affinity and elution assays. In short, this is achieved by including an additional output neuron to the neural network prediction model allowing one prediction for each data type. In this setup, weights are shared between the input and hidden layer for the two input types (binding affinity and eluted ligand), whereas the weights connecting the hidden and output layer are specific for each input type. During neural network training, an example is randomly selected from either data set and submitted to forward and back propagation, according to the NNAlign algorithm. The weight sharing allows information to be transferred between the two data types, and potentially results in a boost in predictive power (for more details on the algorithm, refer to (Jurtz et al., 2017)).

Models were trained and evaluated in a 5 fold cross-validation manner with the same model hyper-parameters that were used for the single data type model. Comparing the performance of the single data type (Table 2), to the multiple data type models for the different data sets (Table 3), a consistent improvement in predictive performance was observed when the two data types were combined. This is the case, in particular, when looking at the PPV performance values. Here, the combined model in all cases has improved performance compared to the single data type model. This is in line to what we have previously observed for MHC class I predictions (Jurtz et al., 2017).

**Table 3.**
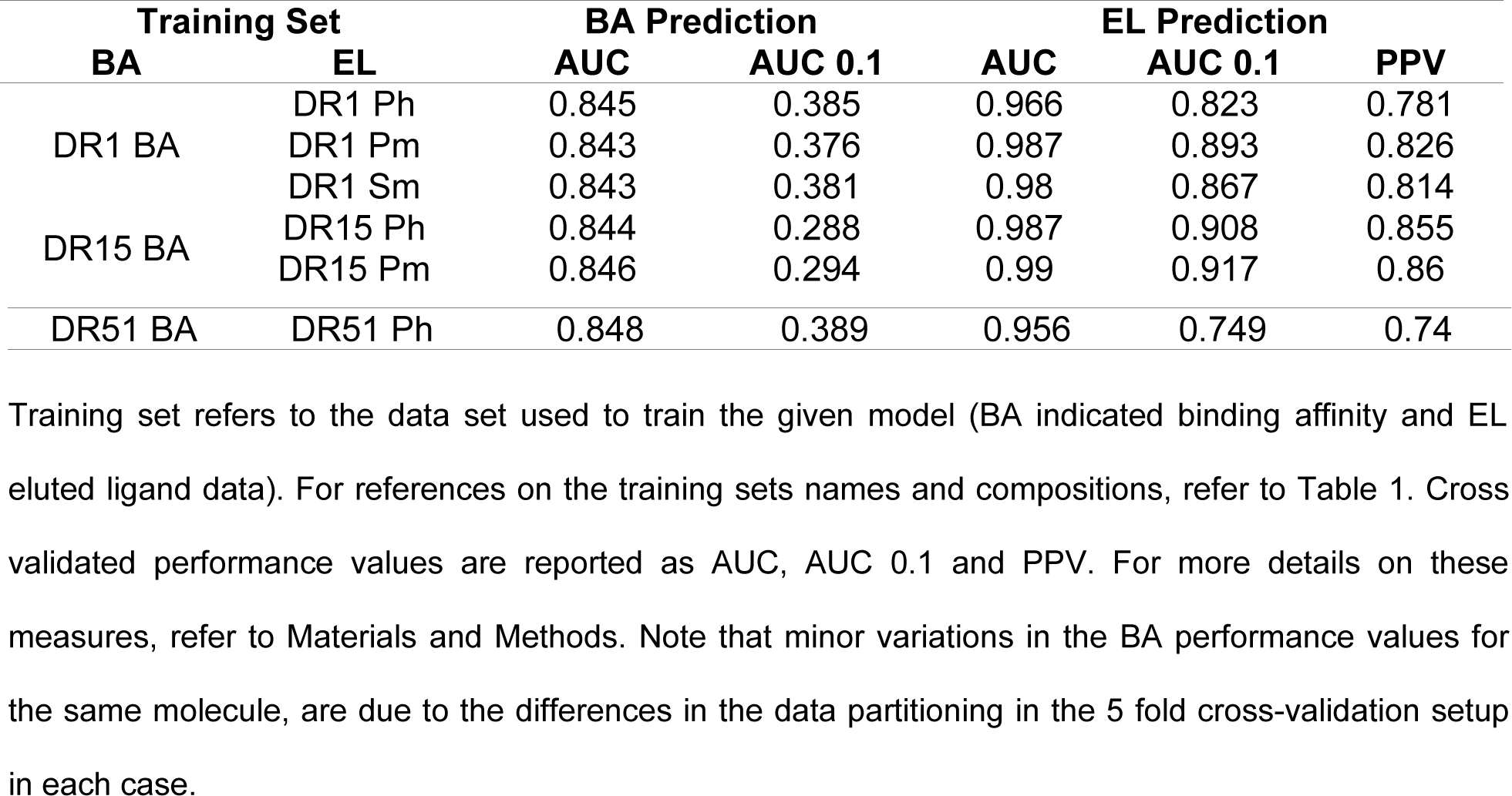
Cross-validation performance for the combined NNAlign models, trained on both Binding Affinity (BA) and Eluted Ligands (EL) data.

Constructing the binding motif captured by the different combined models (see Supplementary Figure 2) confirmed the findings from the single data types model (displayed in Figure 2), with clearly defined and consistent binding motifs in all cases, and with subtle differences in the preferred amino acids at the anchor positions between motifs derived from the binding affinity and eluted ligand output value of the models.

We next turned to the issue of accurately predicting the preferred length of peptides bound to the different HLA-DR molecules. The MS eluted ligand data demonstrated a length preference for the two MHC class II molecules centered on a length around 14-16. Current prediction models such as NetMHCII and NetMHCIIpan are not able to capture this length preference, and have in general a bias of assigning higher prediction values to longer peptides (data not shown). We have earlier demonstrated that including information about the peptide length in a framework integrating MS eluted ligand and peptide binding affinity data, allows the model to capture the length preference of the two data types (Jurtz et al., 2017). Applying a similar approach to the MHC class II data, we obtain the results shown in Figure 3, confirming that also for class II the models are capable of approximating the preferred length preference of each molecule.

**Figure 3.**
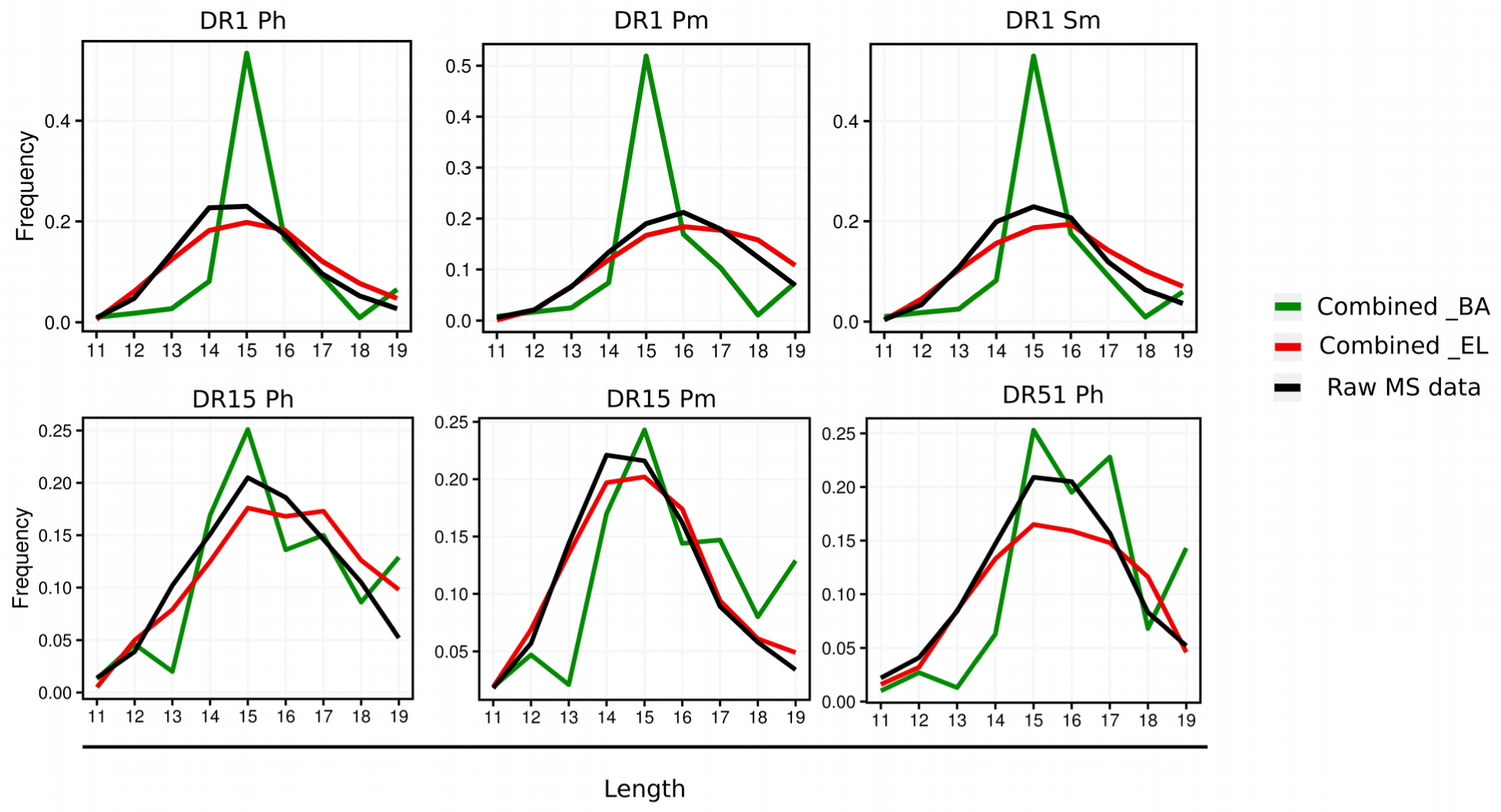
Peptide length preferences learned by the five models trained on Binding Affinity (BA) and Eluted Ligands (EL) combined data. For each model, green traces represent the length histogram of the top 1% scoring predictions for the BA output neuron, on a prediction dataset composed of one million random peptides; red traces refer to the length histogram of the top 0.1% scoring predictions for the EL output neuron, on the same prediction set; black traces indicate the length distribution of the raw MS data.

Lastly, we performed an evaluation across data sets to confirm the robustness of the results obtained, and to reveal any unforeseen signal of performance overfitting. For each data set, we used the two-output model trained above to predict the other ligand data sets of the same allotype. Prior to evaluation, all data with a 9mer overlap between training and evaluation sets were removed. We observed that, in all cases, models trained on a specific data set retained high predictive performance for the prediction of ligands of the same allotype derived from a different experiment (Table 4). These results confirm the high reproducibility of the motifs across different cell lines, as well as the robustness of the prediction models derived from individual data sets.

**Table 4.**
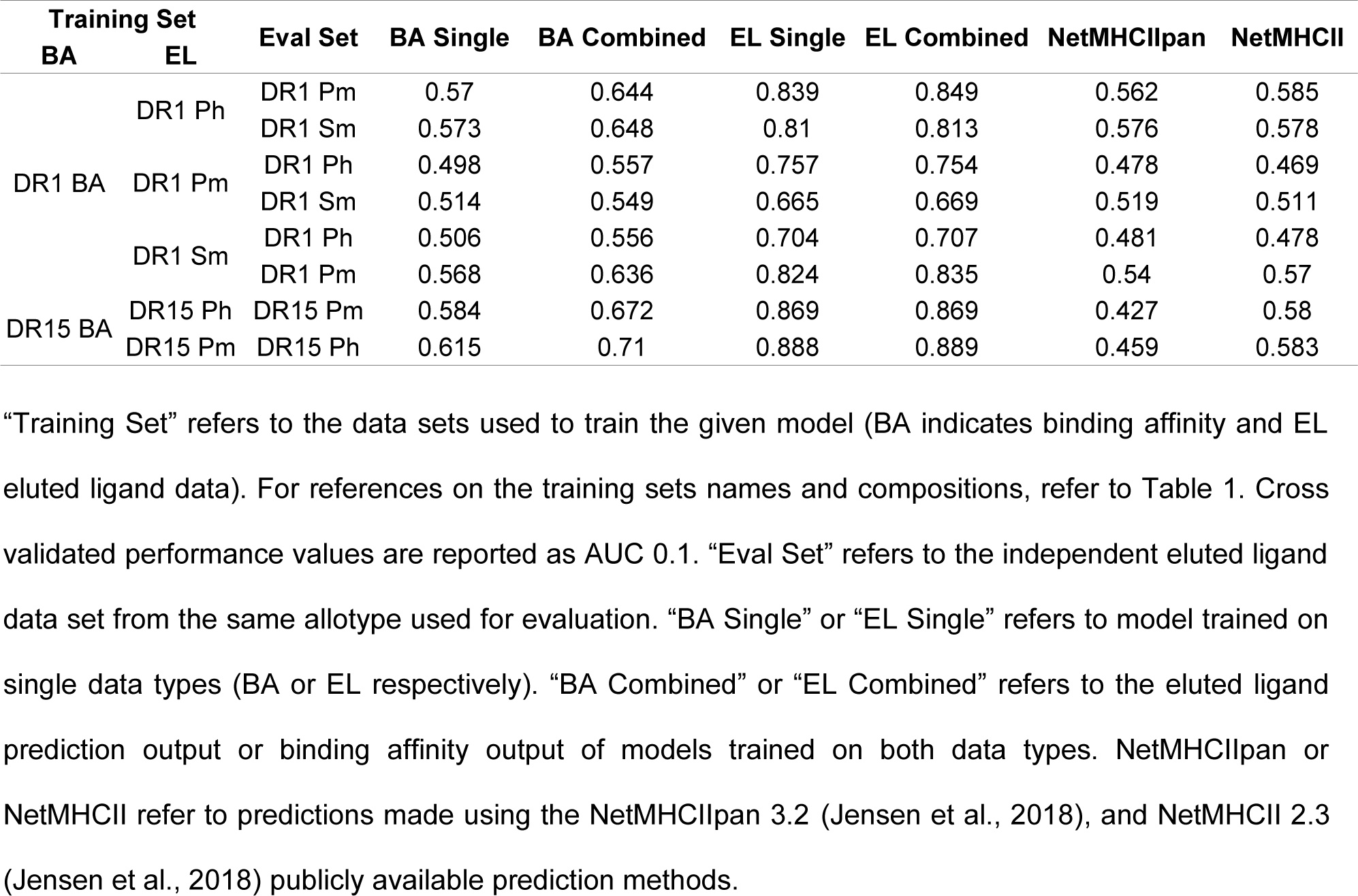
Independent evaluation of eluted ligands dataset in terms of AUC 0.1.

### Signals of Ligand Processing

Having developed improved models for prediction of MHC class II ligand binding, we next analyzed whether the models could be used to identify signals of antigen processing in the MS eluted ligand data sets. We hypothesized that information concerning antigen processing should be present in the regions around the N and C terminus of the ligand. These regions comprise residues that flank the MHC binding core called Peptide Flanking Regions (PFRs), and residues from the ligand source protein sequence located outside the ligand (see lower part of Figure 4 for a schematic overview).

**Figure 4.**
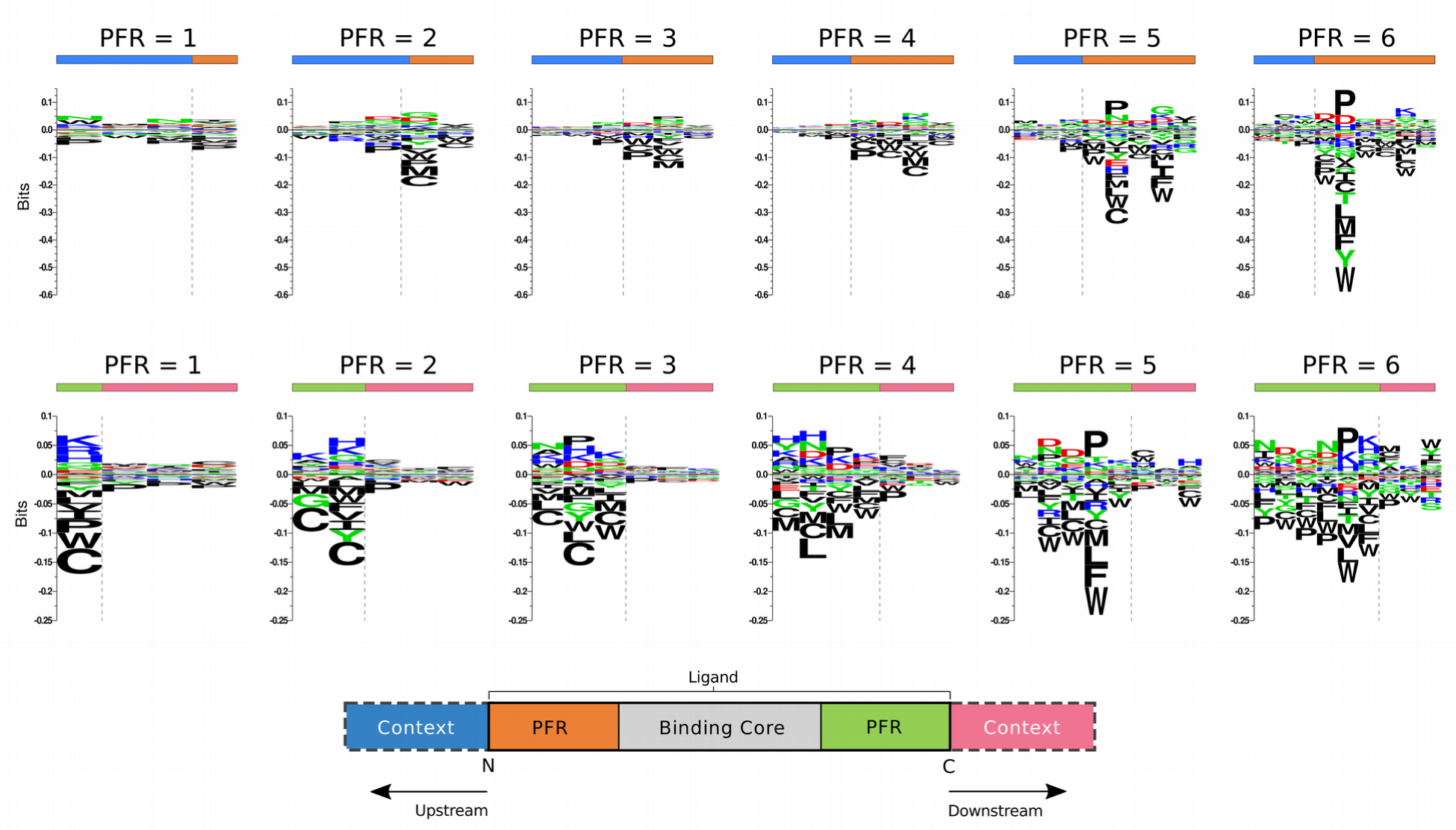
Processing signals found at N and C terminus positions in the DR15 Pm dataset (located at Upstream and Downstream regions, respectively), grouped by Peptide Flanking Regions (PFR) length. For the Upstream part of the ligands (top row), the processing signal is always centered at the N terminal position, extending three positions beyond the cleavage site (Upstream “context”, symbolized as blue bars) and one to six positions towards the binding core, depending on the PFR length (orange bars). For the Downstream region (bottom row), the disposition of elements is mirrored: the proposed processing signal is centered at C terminus, and extends three positions beyond the cleavage site (Downstream “context” region, pink bars) and one to six positions towards the binding core (green bars), depending on the PFR length. Amino acid background frequencies were calculated using the antigenic source protein of all the ligands present in the dataset. Motifs were generated using Seq2logo, as described in Materials and Methods.

We speculate that the signals of antigen processing depend, to some degree, on the length of the PFRs on each side of the binding core. MHC-II ligands are cut and trimmed by exopeptidases, which operate according to specific motifs in prioritizing cleavage sites. However, in the case of short PFRs, the MHC hinders access of the protease to the ligand, hence preventing trimming of the residues in close proximity to the MHC (Larsen, Pedersen, Buus, & Stryhn, 1996; Mouritsen, Meldal, Werdelin, Hansen, & Buus, 1992). For this reason, we expect to observe cleavage motifs only in peptides with sufficiently long PFRs, where the end-of-the-trimming signal is given by the peptide sequence rather than by MHC hindrance. To validate this hypothesis, we identified the PFRs of the ligands in the DR15 Pm EL dataset, as well as three “context” residues found immediately upstream or downstream of the ligand in its source protein. To avoid over-estimation of the performance, the binding core was identified from the cross-validated eluted ligand predictions of the two output model. The ligands were split into groups depending on the length of the C and N terminal PFRs, and sequence logos were generated for each ligand subset using Seq2Logo (Figure 5).

**Figure 5.**
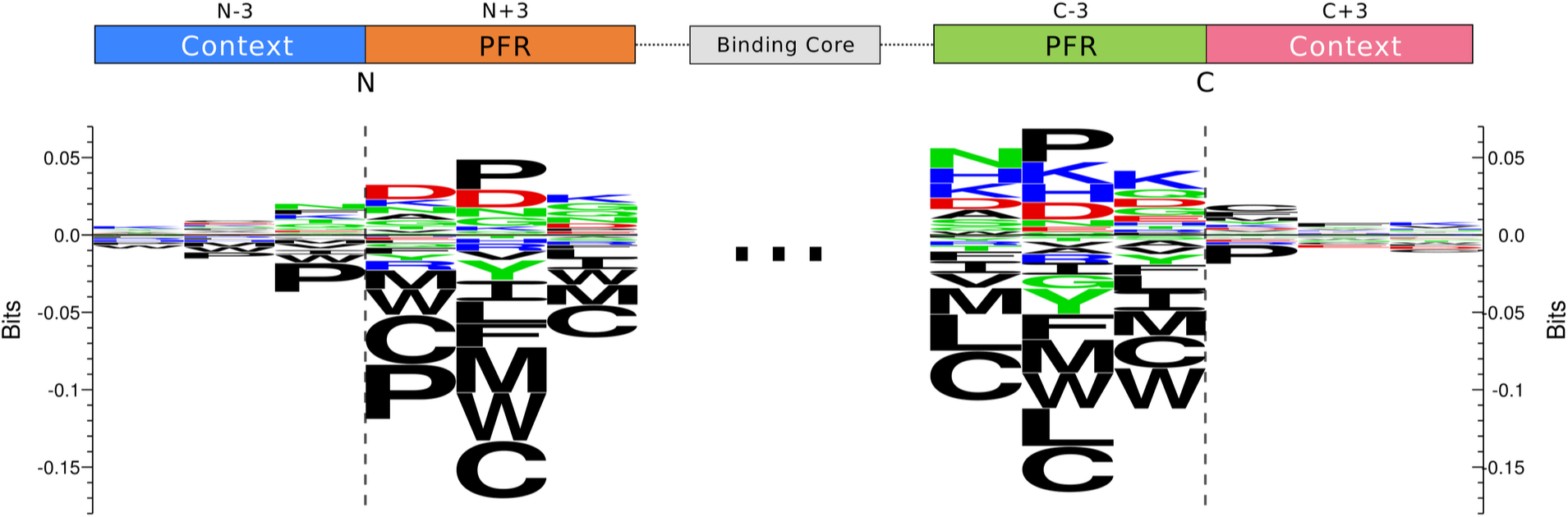
Processing signals located at N and C terminal regions in the DR15 Pm dataset. For each region, all ligands with PFR length lower than 3 were discarded. Then, the logos were constructed as described in the text by selecting the closest three PFR and context residues neighboring the N and C terminus. For additional details on processing signal construction, refer to Figure 4.

The results displayed in Figure 4 clearly confirm the important role of the MHC in shaping the processing signal. For both the N and C terminal datasets, we observe a clear enrichment of proline (P) at the second position from the ligand terminals only for data sets where the PFR is longer than 2 amino acids. This observation is confirmed from the reanalysis of a data set of peptide:HLA-DR complexes from the Protein Data Bank (PDB) previously assembled for benchmarking the accuracy for MHC-II binding core identification (Andreatta et al., 2015). On this PDB dataset, 29% of the entries with a N-terminal PFR longer than 2 amino acids contain a proline at the second position from the N terminal, and 38% of the entries with a C-terminal PFR longer than 2 amino acids contain a proline at the second position from the C terminal (data not shown). On the other hand, none of the bound peptides with N-terminal PFR shorter or equal than 2 amino acids contain a proline at the second position from N-terminal, and only 8% of peptides with C-terminal PFR shorter or equal than 2 amino acids exhibit a proline at the second position from the C-terminal.

To summarize these observations and construct a global motif of the processing signal, we combined the first three C and N terminal residues from all ligands with PFR length larger than two, together with the corresponding three source protein context residues at either C or N terminal side of the ligand. The processing signal at the N and C termini from DR15 Pm is shown in Figure 5; processing motifs for all other data sets can be found in Supplementary Figure 3.

The processing motif confirms the strong preference for proline at the second but last position in the ligand at both N and C termini, as well as a clear signal of depletion of other hydrophobic amino acid types towards the terminals of the ligand. This cysteine depletion in the PFR is likely to be a technological artifact, as cysteines have previously been shown to be underrepresented in MS-derived peptide data sets (Abelin et al., 2017; Bassani-Sternberg et al., 2017). Note also that this depletion is only observed in the PFRs and not in the context residues neighboring the N and C termini. From this figure, it is also clear that processing signals present in the neighborhood (indicated as "context" in Figure 5) of the ligand are very weak. Similar amino acid preferences were obtained in the processing motifs from the other data sets (Supplementary Figure 3).

Next, we investigated to what degree the processing signal was consistently identified in all data sets. To do this, the similarity between any two processing matrices was estimated in terms of the Pearson’s Correlation Coefficient (PCC) between the two vectors of 6*20 elements (6 positions and 20 amino acid propensity scores at each position). The result of this analysis is shown in Figure 6 in terms of a heatmap (the processing matrices from each data set are included in Supplementary Figure 4).

**Figure 6.**
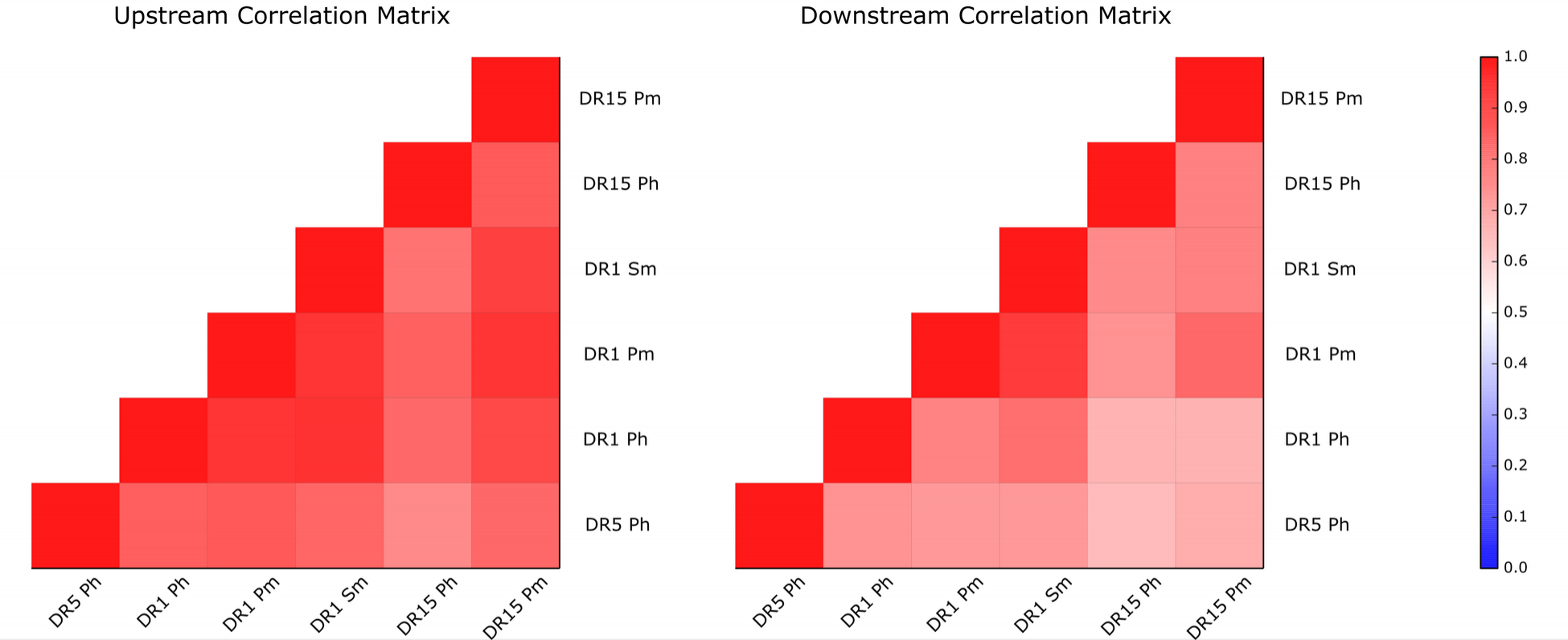
Correlation between processing signals found in the six different datasets employed in this work, for Upstream and Downstream regions. Each matrix entry displays the Pearson Correlation Coefficient (PCC) value of two datasets under study. A PCC value of one corresponds to a maximum correlation, while a PCC value of zero means no correlation. Processing signals used in this figure were generated as explained in Figure 5. All observed PCC values are statistically different from random (p<0.001, exact permutation test).

Figure 6 exhibits a clear positive correlation between the processing motif from all the datasets involved. The mean PCC score for the matrices in Fig. 6 was 0.77 for upstream and 0.73 for downstream, with the lowest PCC=0.59 (for the DR1 Sm and DR1 Ph pair, upstream) and the maximum PCC=0.89 (for DR15 Pm and DR1 Ph, upstream). These results suggest that the processing signals captured are, to a large degree, MHC- and even species-independent: the correlation between the two human and mouse data sets is as high as the correlation between any two data sets within the same species. To ensure that the observed correlation is not related to MS-derived cysteine depletion, we generated the same correlation matrices removing the cysteine contribution and observed no major differences (Supplementary Figure 5). These results thus strongly suggest that the observed signals are related to antigen processing.

### Incorporating ligand processing into a combined predictor

Having identified consistent signals associated with antigen processing, we next investigated whether these signals could be integrated into one model to boost predictive performance. The processing signals were incorporated into the machine-learning framework by complementing the encoding of each ligand with the 3 N terminal context, 3 N terminal peptide, 3 C terminal context, and 3 C terminal peptide residues (see figure 5). For peptide binding affinity data, the context information was presented to the neural networks with three wildcard amino acids “XXX”, corresponding to a vector of zeros. Two models were trained for each one of the allotypes considered in this work: one model including and one excluding the context information, both allowing integration of binding affinity and eluted ligand data. Prior to training, the complete set of data (binding affinity and eluted ligands for all three MHC-II molecules) was split into 5 partitions using the common-motif approach as described in Materials and Methods. All model hyper-parameters were identical to the ones used earlier. The result of this benchmark is shown in Table 5, and confirms that the inclusion of context leads to a consistently improved predictive power of the models for all three data sets.

**Table 5.** Cross-validation performance for combined NNAlign models trained on single-allele datasets, with and without context information.

| **Allele** | **Without context** |  | **With context** |  |
| --- | --- | --- | --- | --- |
|  | **AUC 0.1** | **PPV** | **AUC 0.1** | **PPV** |
| DRB1*01:01 | 0.874 | 0.824 | 0.893 | 0.839 |
| DRB1*15:01 | 0.931 | 0.875 | 0.947 | 0.892 |
| DRB5*01:01 | 0.805 | 0.76 | 0.818 | 0.782 |

“Allele” refers to the combination of all datasets for that given allele used to train the model. Cross validated performance values are reported as AUC 0.1 and PPV. For more details on these measures, refer to Materials and Methods.

As an example of the processing signal captured by a model trained including context information, we constructed sequence motifs of the top 1% highest scoring peptides from a list of 1 million random natural peptides of length 10-25 and their context, for a combined model trained on the DR15 Pm dataset (Supplementary Figure 5). As expected, the motif contained within the N and C terminal peptide flanks and context is close to identical to the motif described in Figure 5.

### T cell epitope prediction using the combined models

Having observed how prediction of naturally processed MHC ligands benefited from implementing ligand context features, we next wanted to evaluate if a similar gain could be observed when predicting T cell epitopes. We downloaded all available epitopes of length 14 to 19 (included) from the IEDB, for the molecules DRB1*01:01, DRB1*15:01 and DRB5*01:01. After filtering out entries with post translational modifications and entries lacking information about the source protein IDs, a total of 557, 411 and 114 epitopes remained for the three DR molecules, respectively. First, we evaluated this panel of epitopes in a conventional way: digesting the epitope source protein into overlapping peptides with the length of the epitope; predicting the peptides using the different models; and calculating the AUC (area under the receiver operator curve) per source protein-epitope pair, taking peptides identical to the epitope as positives and all other peptides in the source protein as negatives. We excluded from the evaluation datasets negative peptides that shared a common motif of 9 amino acids with the epitope. Four methods were included in this benchmark: EL (the eluted ligand prediction value from the model trained on the combined data without context information), EL+context (the eluted ligand prediction value from the model trained on the combined data including context signals), NetMHCII (version 2.3) and NetMHCIIpan (version 3.2). This analysis shows, in line with what we observed earlier for the eluted ligand benchmarks, a consistent improved performance of the EL model compared to both NetMHCII and NetMHCIIpan (Figure 7a).

**Figure 7.**
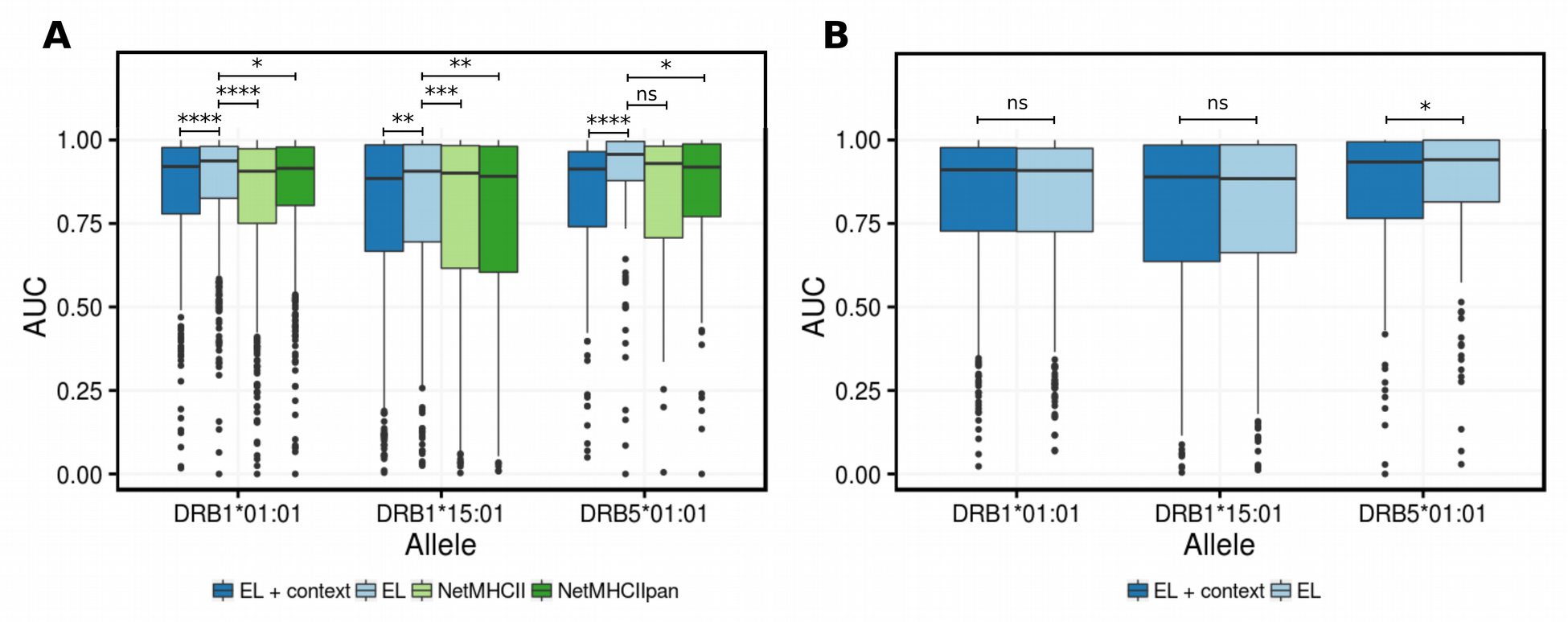
Predictive performance on a panel of CD4+ T-cell epitopes. The boxplots represent the distribution of AUC values over all epitope evaluation datasets restricted to a given allele comparing the different models. Middle lines in boxes correspond to median values. The height of the box represents 50% of the data. Whiskers represent 1.5 quartile range (QR) of data and dots represent outliers of 1.5 of QR. P-significance is calculated from Wilcoxon test. ns (P > 0.05), * (P ≤ 0.05), ** (P ≤ 0.01), *** (P ≤ 0.001), **** (P ≤ 0.0001). In both benchmarks, an AUC value was calculated for each epitope/source protein pair by considering peptides identical to the epitope as positives and all other peptides as negatives excluding peptides with an overlap of at least 9 amino acids to the epitope. **A)** Comparison of the combined models developed in this study with context information (EL + context) and without context (EL) to current state of the art prediction methods trained on binding affinity data only (NetMHCII-2.3 and NetMHCIIpan-3.2). **B)** Comparison of EL + context and EL in a benchmark where the epitope evaluation set was constructed using the evaluation strategy accounting for ligand preference described in the text.

The benchmark however also demonstrates a substantial drop in predictive power of the EL model when incorporating the context processing signal (EL+context). This drop is however expected since the mapped T cell epitope boundaries are not a product of natural antigen processing and presentation, but rather result from screening of overlapping peptides from a candidate antigen, or by peptides synthesized based on the results of MHC peptide binding predictions and/or *in vitro* binding assays. As a consequence, the N and C terminal boundaries of such epitope peptides do not necessarily contain the processing signal obtained from naturally processed ligands. However, given that the epitope was demonstrated to bind to the T cell originally induced towards a naturally processed ligand, we can assume that the sequence of the validated epitope and the original (but unknown to us) naturally processed ligand share an overlap at least corresponding to the MHC-II binding core of the validated epitope. Following this reasoning, we redefined the epitope benchmark as follows. First, we predicted a score for all 13-21mer peptides within a given source protein using the EL or EL+context models. Next, we digested the source protein into overlapping peptides of the length of the epitope, and assigned a score to each of these peptides corresponding to the average prediction score of all 13-21mer peptides sharing a 9mer or more overlap with the given peptide (models where the max score was assigned were also considered, but gave consistently lower predictive performance, data not shown). Finally, we calculated as before an AUC value for the epitope-source protein pair taking peptides equal to the epitope as positives and all other peptides as negatives excluding from the evaluation set negative peptides sharing a common motif of 9 amino acids with the epitope. The benchmark shows a comparable performance of the EL+context method vs EL method for the alleles analyzed in the study (Figure 7b). Possible reasons for this lack of improved performance of the EL+context model are discussed below.

## Discussion and Conclusions

Peptide binding to MHC II is arguably the most selective step in antigen presentation to CD4+ T cells. The ability to measure (and predict) specific CD4+ responses is crucial for the understanding of pathological events, such as infection by pathogens or cancerous transformations. Recent studies have also highlighted a potential role for CD4+ T cells for the development of cancer immunotherapies (Kreiter et al., 2015; Tran et al., 2014; Zanetti, 2015). Characterizing peptide:MHC-II binding events has been a focal point of research over the last decades. Large efforts have been dedicated in conducting high-throughput, *in vitro* measurements of peptide MHC II interactions (Justesen, Harndahl, Lamberth, Nielsen, & Buus, 2009; J. Sidney et al., 2001; John Sidney et al., 2013), and these data have been used to develop methods capable of accurately predicting the interaction of peptides to MHC II molecules from the sequence alone (Andreatta et al., 2015; Nielsen & Andreatta, 2017; Nielsen, Lundegaard, & Lund, 2007; Singh & Raghava, 2001). While these approaches have proven highly successful as guides in the search for CD4 epitopes (Gfeller, Bassani-Sternberg, Schmidt, & Luescher, 2016; Nielsen, Lund, Buus, & Lundegaard, 2010), a general conclusion from these studies is that MHC II *in vitro* binding affinity (whether measured or predicted) is a relatively poor correlate of immunogenicity (Backert & Kohlbacher, 2015). In other words, peptide binding affinity to MHC II is a necessary but not sufficient criterion for peptide immunogenicity. The same situation holds for MHC class I presented epitopes. Here, however, peptide binding to MHC I is a very strong correlate to peptide immunogenicity, and can be used to discard the vast majority (99%) of the irrelevant peptide space while maintaining an extremely high (>95%) sensitivity for epitope identification (Jurtz et al., 2017). For MHC II, recent studies suggest that the corresponding numbers fall in the range 80% specificity and 50% sensitivity (Jensen et al., 2018). For these reasons, we suggest that other features than MHC II in-vitro binding affinity may be critical for MHC II antigen presentation. Based on six MS MHC II eluted ligand data sets, we have here attempted to address and quantify this statement.

Firstly, we have demonstrated that the MS MHC II eluted ligand data sets employed in this work (generated by state-of-the-art technologies and laboratories) are of very high quality, with low noise levels and allowing very precise determination of MHC II binding motifs. Overall, the obtained binding motifs show overlap with the motifs identified from in-vitro binding affinity data, with subtle differences at well-defined anchor positions.

Secondly, we demonstrated that high accuracy prediction models for peptide MHC II interaction can be constructed from the MS-derived MHC II eluted ligand data; that the accuracy of these models can be improved by training models integrating information from both binding affinity and eluted ligand data sets; and that these improved models can be used to identify both eluted ligands and T cell epitopes in independent data sets at an unprecedented level of accuracy. This observation strongly suggests that eluted ligand data contain information about the MHC peptide interaction that is not contained within *in vitro* binding affinity data. This notion is further supported by the subtle differences observed in the binding motifs derived from eluted ligand and *in vitro* binding affinity data. Similar observations have been made for MHC class I (Abelin et al., 2017; Jurtz et al., 2017). We at this point have no evidence for the source of these differences, but a natural hypothesis would be that they are imposed by the presence of the molecular chaperones (such as HLA-DM) present in the eluted ligand but absent from *in vitro* binding assays. An alternative explanation could be that the eluted peptide ligands reflect peptide-MHC class II stability rather than affinity; something that would imply that stability is a better correlate of immunogenicity than affinity (Lazarski et al., 2005).

Thirdly, we analyzed signals potentially associated with antigen processing. We postulate that such processing signal should be influenced by the relative location of the peptide binding core compared to the N and C terminal of the given ligand. This is because the MHC II molecule may hinder the access of the protease, thus preventing trimming of the residues in close proximity to the MHC (Larsen et al., 1996). Investigating the data confirmed this hypothesis, and a relatively weak but consistent processing signal (with a preference for Prolines at the second amino acid position from the N and C terminal of the ligand), was observed for ligands where the length of the region flanking the binding core was three amino acids or more. This observation was found consistently in all data sets independent of MHC II restriction and host species (human or mouse).

Lastly, we integrated this information associated with antigen processing into a machine-learning framework, and demonstrated a consistently improved predictive performance not only in terms of cross-validation but also when applied to independent evaluation data sets covering naturally processed MHC eluted ligands. However, we do not observe an improvement of the extended model for prediction of validated T cell epitopes. There are several possible reasons for this. In the first place, it is possible that epitope data have a bias towards current MHC class II binding prediction and/or in vitro binding assay methods, since researchers could use these tools to select which peptides to include in a T cell epitope screening or to define the MHC restriction element for a given positive epitope. Secondly, we have attempted a very simple strategy to assign a prediction score to each epitope. It might be that the conclusion is altered if alternative, more sophisticated mapping strategies were used. Finally, the reason might be biological: the antigen processing pathways predominantly utilized in cell lines used for ligand elution experiments which lead to the motifs we identified might not be the only ones generating T cell epitopes *in vivo*, where e.g. cross-presentation might play a role. Future work is needed to disentangle this question.

In conclusion, we have demonstrated how integrating MHC class II *in vitro* binding and MS eluted ligand data can boost the predictive performance for both binding affinity, eluted ligand, and T cell epitope predictions. To the best of our knowledge, we have also demonstrated for the first time how MHC II eluted ligand data can be used to extract signals of antigen processing, and how these signals can be integrated into a model with improved predictive performance.

Our work is limited to three HLA-DR molecules, but the framework can be readily extended to additional molecules, once sufficient data become available. Also, it may become achievable to construct a pan-specific predictor as has been shown earlier for MHC class I (Jurtz et al., 2017), enabling predictions for any MHC molecule of known sequence.

## Declarations

### Ethics approval and consent to participate

Not applicable

### Availability of data and materials

The eluted ligand datasets analyzed during the current study are available in the IEDB repository, http://www.iedb.org/reference/1031891 and http://www.iedb.org/reference/1030084.

### Authors’ contributions

MN designed the study. CB, BA, MA and MN performed the experiments and statistical analysis. CB, BA, SP, AS, BP, SB, MA, and MN analyzed and interpreted the data, and wrote the paper. All authors read and approved the final manuscript.

## List of Abbreviations

MHC-II: Major histocompatibility complex class-II
pMHCII: peptide-MHC-II complexes
PFRs: Peptide flanking regions
MS: Mass spectrometry
PSSM: Position specific scoring matrix
KLD: Kullback-Leibler distance
PCC: Pearson correlation coefficient
BA: Binding Affinity data
EL: Eluted Ligands data
AUC: Area Under the ROC Curve
AUC 0.1: Area Under the ROC curve integrated up to False Positive Rate of 10%
PPV: Positive Predictive Value

